# Sensory Circuit Dynamics in a Mouse Model of Epileptic Encephalopathy

**DOI:** 10.1101/2025.11.22.689933

**Authors:** Dymphie Suchanek, Anthony Williams, Tianyu Cao, William D. Todd, Qian-Quan Sun

## Abstract

Epileptic encephalopathies (EEs) are severe developmental disorders with abnormal EEG activity that worsens neurodevelopmental deficits, yet underlying sensory and cognitive impairment mechanisms are unclear. We studied cortical circuit dynamics in Ank3-1b^-/-^ mice, an EE model with parvalbumin interneuron dysfunction, using laminar electrophysiology, spike-field coherence (SFC), single-unit analyses, and behavioral assays. Ank3-1b^-/-^ mice showed disrupted excitatory-inhibitory (E-I) balance, with reduced sink/source ratios (p = 0.0279) and elevated net currents (AVREC; p < 0.01), indicating hyperexcitability. In awake states, sensory-evoked intra-columnar connectivity was impaired, with lower Pearson correlations (p < 0.001) and increased low-frequency (3–30 Hz) SFC (p < 0.001), but intact gamma-band coherence, suggesting aberrant synchronization. Inhibitory neuron latencies were delayed in infragranular layers under anesthesia (p < 0.001) and supragranular layers when awake (p = 0.005), implying thalamocortical deficits. Despite preserved sensory adaptation and recognition memory, Ank3-1b^-/-^ mice exhibited heightened anxiety (p < 0.05) and variable circadian rhythms, indicating selective affective and regulatory deficits. These results show that Ank3-1b loss disrupts cortical E-I balance and synchrony, delaying sensory processing, while compensatory mechanisms maintain critical sensory functions. This study links ANK3 mutations to layer-specific circuit dysfunction, offering insights into EE pathophysiology and sensory deficits.

## Introduction

Epileptic encephalopathies (EEs) are severe developmental and epileptic disorders characterized by electroclinical features that exacerbate neurodevelopmental impairment (McTague, Howell et al. 2016, Shao and Stafstrom 2016). Their EEG patterns are highly heterogeneous, often displaying sustained, rhythmic, and unreactive abnormalities. While some syndromes exhibit bilateral synchronous discharges (e.g., Lennox-Gastaut syndrome with slow spike-and-wave complexes(Nariai 2025)), others show focal, multifocal, or migrating epileptiform activity^13^ (e.g., Dravet syndrome or early infantile epileptic encephalopathy with migrating focal seizures). Spatiotemporal evolution may occur during seizures, across developmental stages (e.g., Ohtahara syndrome progressing to West syndrome), or in specific sleep-activated states (e.g., electrical status epilepticus during sleep [ESES] in Landau-Kleffner syndrome). Dynamic EEG patterns and frequent drug-resistant seizures in EEs worsen cognitive and behavioral decline (McTague, Howell et al. 2016, Shao and Stafstrom 2016). However, the relationship between EEs and cognitive deficits, as well as the underlying mechanisms, remains poorly understood.

Sensory processing abnormalities, such as hypersensitivity to stimuli in Dravet syndrome (DS) (Zuberi, Wirrell et al. 2024) or integration deficits in Lennox-Gastaut syndrome (LGS) (Arzimanoglou, French et al. 2009), are common but understudied. Cognitive impairments, including intellectual disability and memory deficits, are hallmarks of EEs(Berg, Berkovic et al. 2010), and their severity correlates with the burden of interictal epileptiform discharges (Shao and Stafstrom 2016). Preclinical studies suggest these deficits arise from disrupted cortical circuits, particularly involving parvalbumin (PV) interneuron dysfunction(Lopez, Wang et al. 2016, Tatsukawa, Ogiwara et al. 2018). For example, Scn1a haploinsufficient mice (a DS model) exhibit somatosensory deficits and impaired hippocampal plasticity (Han, Yu et al. 2012, Favero, Sotuyo et al. 2018), linking inhibitory dysfunction to sensory and cognitive phenotypes. However, the role of cortical network dynamics, including columnar processing of sensory inputs and their modulation by EEs, remains largely unexplored.

The ANK3 gene, encoding ankyrin-G (AnkG), regulates neuronal excitability by clustering NaV and KCNQ2/3 channels at axonal initial segments(Pan, Kao et al. 2006, Rasband 2010). ANK3 variants are strongly implicated in bipolar disorder (Ferreira, O’Donovan et al. 2008, Muhleisen, Leber et al. 2014) and epilepsy(Lopez, Wang et al. 2017), with six of its ligand genes (SCN1A, SCN2A, SCN8A, SCN1B, KCNQ2, KCNQ3) directly linked to EEs(Noebels 2015, McTague, Howell et al. 2016). This dual role positions ANK3 at the nexus of epilepsy and psychiatric comorbidity, a relationship poorly addressed by current therapies(Geddes and Miklowitz 2013). In Ank3-1b^-/-^ mice, deletion of the AnkG exon 1b isoform selectively reduces NaV channel density at PV interneuron axonal initial segments, causing hypoexcitability, seizures, and sudden death (Lopez, Wang et al. 2016).Chronic EEG recordings in these mice reveal continuous spike-wave discharges during slow-wave sleep, mimicking human ESES, alongside progressive hippocampal involvement and interictal spikes with fast ripples (250–500 Hz) (Lopez, Wang et al. 2016). These features mirror the aberrant nested oscillations hypothesized to drive EE pathophysiology (Lenck-Santini 2017). However, sensory and cognitive deficits in this model, and their circuit dynamics during sensory processing, remain uncharacterized.

Here, we investigate sensory processing and cognitive impairments in Ank3-1b^-/-^ mice using *in vivo* laminar electrophysiology, spike-field coherence (SFC), single-unit dynamics, and behavioral assays. We hypothesize that PV interneuron dysfunction destabilizes cortical excitation-inhibition (E-I) balance and disrupts network synchrony, leading to sensory hypersensitivity and anxiety-like behaviors. By linking cellular and network excitability deficits to altered oscillatory dynamics and behavior, our findings aim to identify circuit-level mechanisms and therapeutic targets for EEs and their comorbidities.

## Results

### 1. Cortical columnar excitation and inhibition balance

To determine if Ank3-1b disruption models a specific epileptic encephalopathy (EE), we characterized sleep neurophysiology. We found that all adult Ank3-1b^-/-^ mice (n = 15/15) exhibited a severe, continuous spike-wave during sleep (CSWS) phenotype(Sun, Zhou et al. 2016) (Fig 1C), also known as electrical status epilepticus in sleep (ESES)(Brazzo, Pera et al. 2012, Sun, Zhou et al. 2016, Chapman, Haubenberger et al. 2024). This was characterized by abundant, high-amplitude spike-wave discharges occupying >80% of NREM sleep epochs. These electrographic seizures were associated with pathological high-frequency oscillations (HFOs) and increased neuronal firing in the somatosensory cortex area (Fig 1C), indicating the ssomatosensory cortex as a primary site of epileptogenesis(Williams and Sun 2019, Shi, Shaw et al. 2024). The universal presence of this CSWS/ESES syndrome establishes the adult Ank3-1b mutant mouse as a robust model for investigating this devastating epileptic encephalopathy.

**Figure 1.**
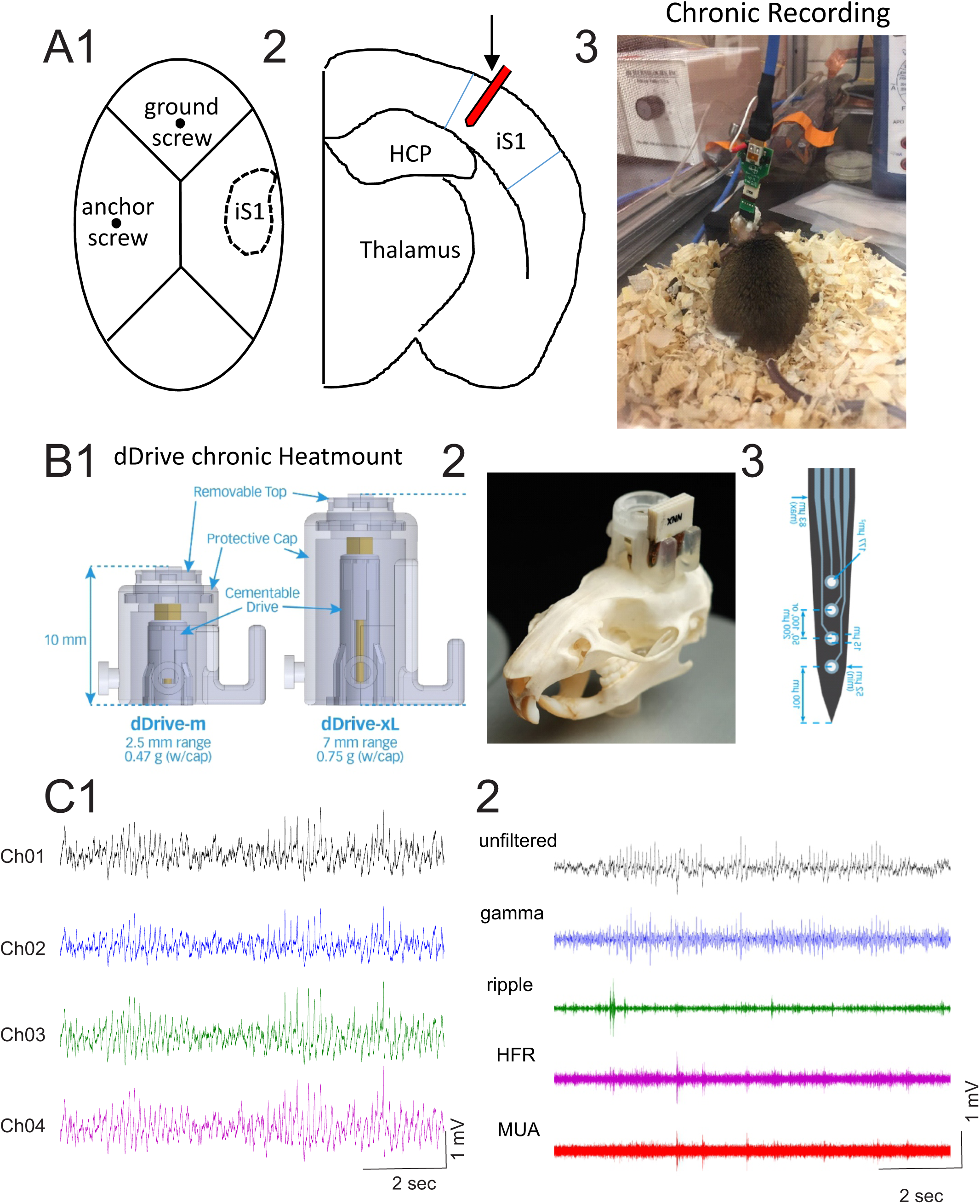
Adult Ank3-1b mutant mice model epileptic encephalopathy with CSWS during sleep. (A) Schematic (A1, A2) and photo (A3) of chronic electrode implantation in S1. (B) Illustration of a “quatrode” microdrive. (C) Representative sleep recordings showing: (C1) raw LFP with spike-wave discharges (SWDs), (C2) band-pass filtered (80-200 Hz) trace revealing high-frequency oscillations (HFOs), and (C3) concurrent single-unit raster plots from two neurons during SWDs.

Linear electrodes (16 channels) were implanted across cortical columns in the barrel cortex to record air-puff-induced cortical responses under either isoflurane anesthesia or fully awake conditions (Fig. 2). Vibrissa-evoked cortical field potentials are generated by a basic circuit of sequential supragranular-to-infragranular pyramidal cell activation, producing fast depolarizing and slow repolarizing current sequences(Di, Baumgartner and Barth 1990). The Vibrissa-evoked cortical field potentials was found to unfold in three sequential phases: initial thalamic activation of layers IV (Fig 3A2), followed by intracortical supragranular activation. These responses represent precise spatiotemporal sequence of microcircuit recruitment that generates cortical field potentials(Di, Baumgartner and Barth 1990, Jellema, Brunia and Wadman 2004).

**Figure 2.**
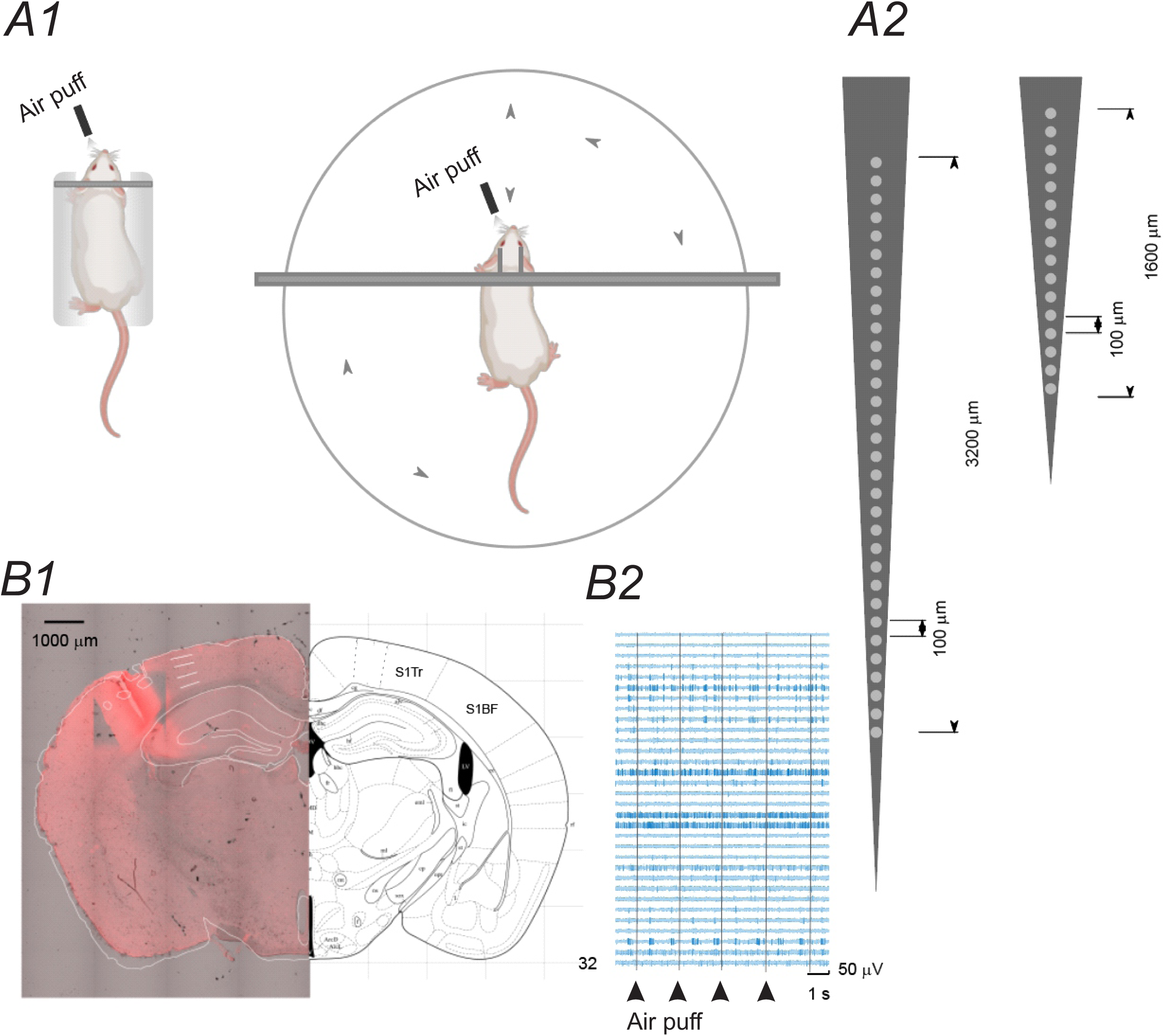
In vivo laminar recording setup for assessing sensory-evoked cortical dynamics. (A) Schematic of the head-fixed, freely moving recording configuration within a Neurotar Mobile HomeCage™. An air puff was delivered to the whisker pad while laminar local field potentials (LFPs) and single-unit activity were recorded from the barrel cortex using a chronically implanted two different 16-channel silicone probes (NeuroNexus Technologies). (B1) Histological verification of the recording site. Left: Photomicrograph of a coronal section showing the electrode track (arrow). (B2) An example of MUAs (multi-unit activities) evoked by air-puff stimulations.

**Figure 3.**
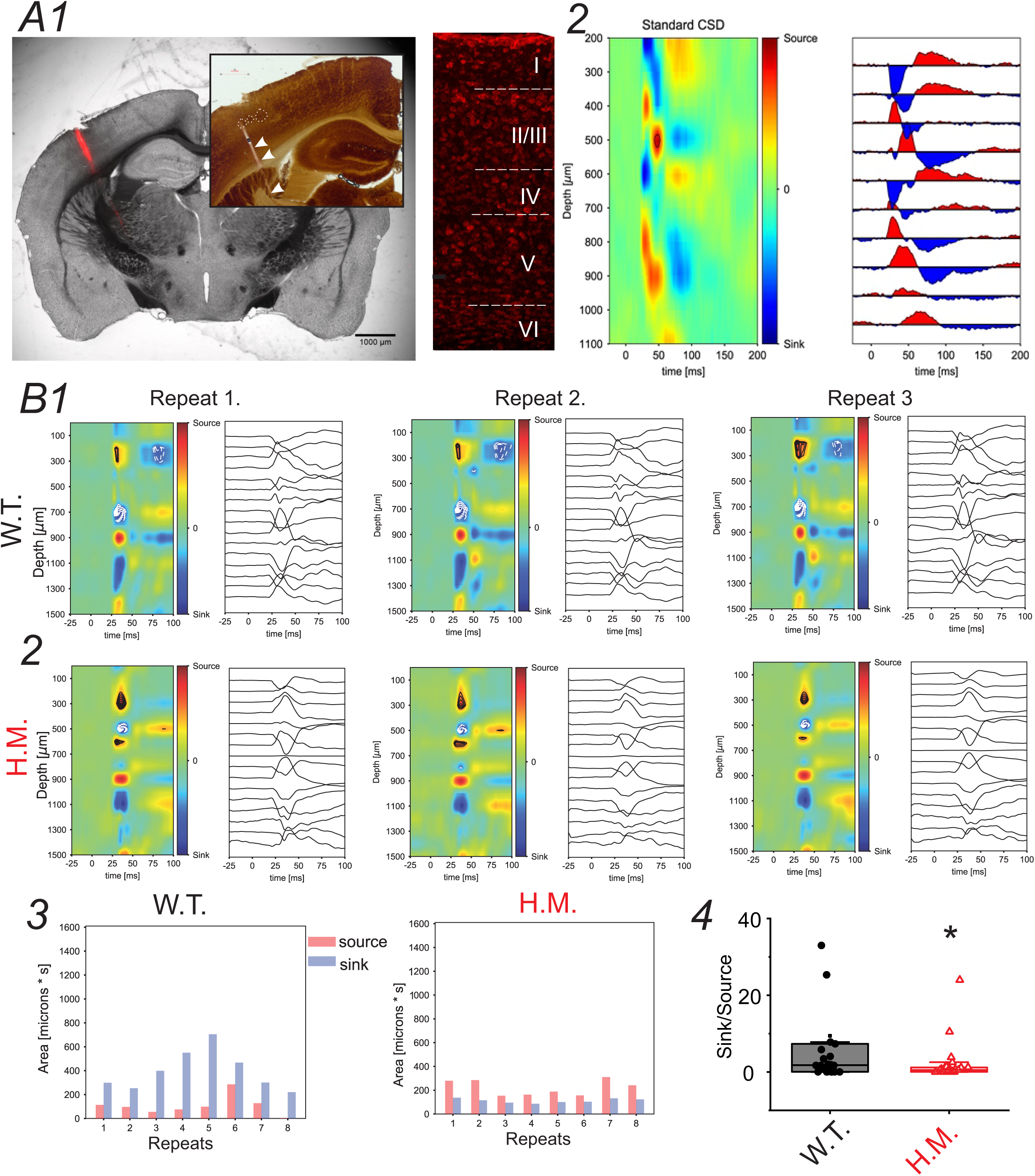
Ank3-1b mutation disrupts laminar excitation-inhibition balance in the barrel cortex. (A) Representative current source density (CSD) plots in response to whisker stimulation in awake, behaving mice. Sinks (net excitatory currents, red) and sources (net inhibitory currents, blue) are shown across cortical depth and time. (B1) CSD from a wild-type (WT) mouse. (B2) CSD from an Ank3-1b^⁻/⁻^ (H.M.) mouse. (B3) Quantification of the total area of sensory-evoked sinks and sources over 500 ms post-stimulus from two representative animals. (C) Scatter and box plot comparing the sink/source ratio between genotypes (p = 0.0279, unpaired t-test, df=49).

Current source density (CSD) analysis(Nicholson and Freeman 1975) revealed distinct alterations in sensory-evoked activity between Ank3-1b^-/-^ and wild-type (WT) mice. Quantitative comparisons of sink-source dynamics demonstrated a significantly lower sink/source ratio in Ank3-1b^-/-^ mice (p = 0.0279, unpaired t-test, df = 49; Fig. 2B4), driven by a marked decrease in total sink area (Ank3-1b^-/-^: 167.0 ± 29.8 vs. WT: 289.5 ± 51.3 µm²·s, p = 0.034, df = 49) and opposing changes in source magnitudes (Ank3-1b^-/-^: 454.8 ± 69.8 vs. WT: 234.3 ± 60.6 µm²·s, p = 0.026, df = 49, Fig 3B3). This imbalance suggests disrupted coordination of excitatory and inhibitory currents in Ank3-1b^-/-^ mice. Furthermore, the average rectified current (AVREC), a measure of net transmembrane current flow, was significantly larger in Ank3-1b^-/-^ mice (p < 0.01; Fig. 4), peaking at 0.0023 ± 0.0025 µV/mm² in Ank3-1b^-/-^ mice compared to 0.0008 ± 0.0015 µV/mm² in WT under fully awake conditions (Fig. 4D, p < 0.0001), but not under anesthetized conditions (Fig. 4C, p = 0.327). The amplified AVREC in Ank3-1b^-/-^ mice aligns with hyperexcitable cortical states, likely reflecting impaired parvalbumin interneuron-mediated inhibition. Together, these findings highlight aberrant sensory-evoked laminar processing in Ank3-1b^-/-^ mice, characterized by exaggerated excitatory sink activity and diminished inhibitory regulation, which may underlie the sensory and cognitive deficits observed in this model of EE.

**Figure 4.**
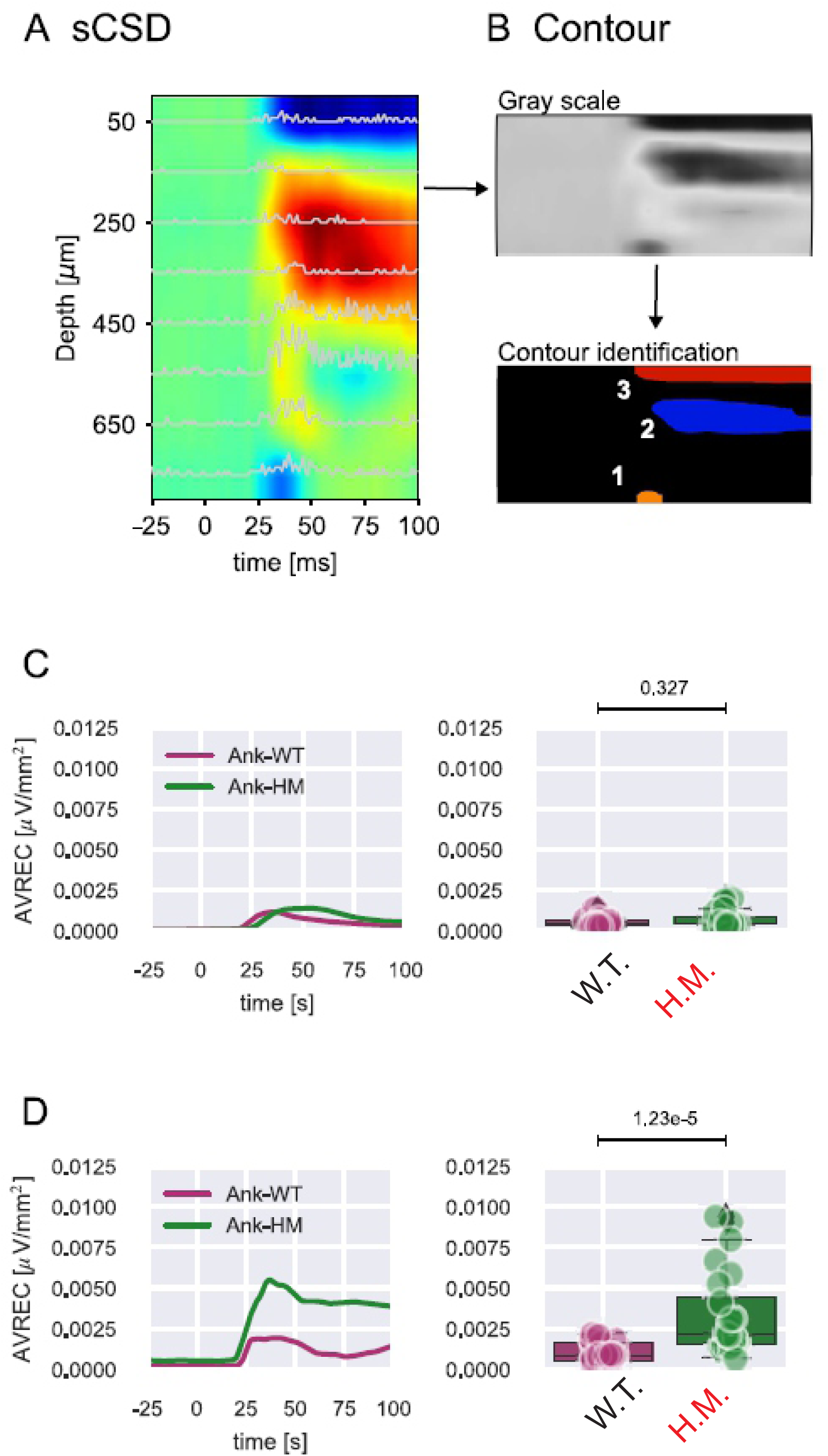
Enhanced net cortical excitability in awake Ank3-1b^⁻/⁻^ mice revealed by Average Rectified Current (AVREC) analysis. (A-B) Conversion of CSD maps to AVREC. (A) Representative CSD plot from a WT mouse. (B) The same CSD data converted to a grayscale absolute-value map with automatically identified contours (colored lines). Schematic illustrating the AVREC calculation by rectifying and averaging the CSD signal across all cortical depths. (C) Peak AVREC magnitude under isoflurane anesthesia (p = 0.327, unpaired t-test). Traces represent mean ± SEM. (D) Peak AVREC magnitude in the awake, behaving state (p < 0.0001, Mann-Whitney-U-test).

### 2. Intra-columnar network synchronizations

Cross-correlation analysis of unfiltered intra-columnar cortical network activity revealed pronounced deficits in functional connectivity in Ank3-1b^-/-^ mice compared to WT controls. Pearson correlation coefficients, quantifying pairwise synchrony between cortical layers during whisker-evoked responses, were significantly reduced for immediate thalamocortical responses in Ank3-1b^-/-^ mice (Ank3-1b^-/-^: 0.65 ± 0.21 vs. WT: 0.76 ± 0.15; p < 0.001, mixed-effects ANOVA; Fig. 5B3). This reduction was observed only in awake conditions, not under anesthesia (Fig. 5B1), and was most pronounced in supragranular layers (II/III) and thalamocortical recipient zones (layer IV). Hierarchical clustering of cross-correlation matrices further demonstrated impaired network organization in awake Ank3-1b^-/-^ mice, as evidenced by a lower cophenetic correlation coefficient (Ank3-1b^-/-^: 0.78 ± 0.05 vs. WT: 0.91 ± 0.09; p < 0.001, Mann-Whitney U-test; Fig. 5B4) compared to anesthetized conditions (Fig. 5B2), indicating disrupted modularity of cortical microcircuits during sensory processing in fully awake states.

**Figure 5.**
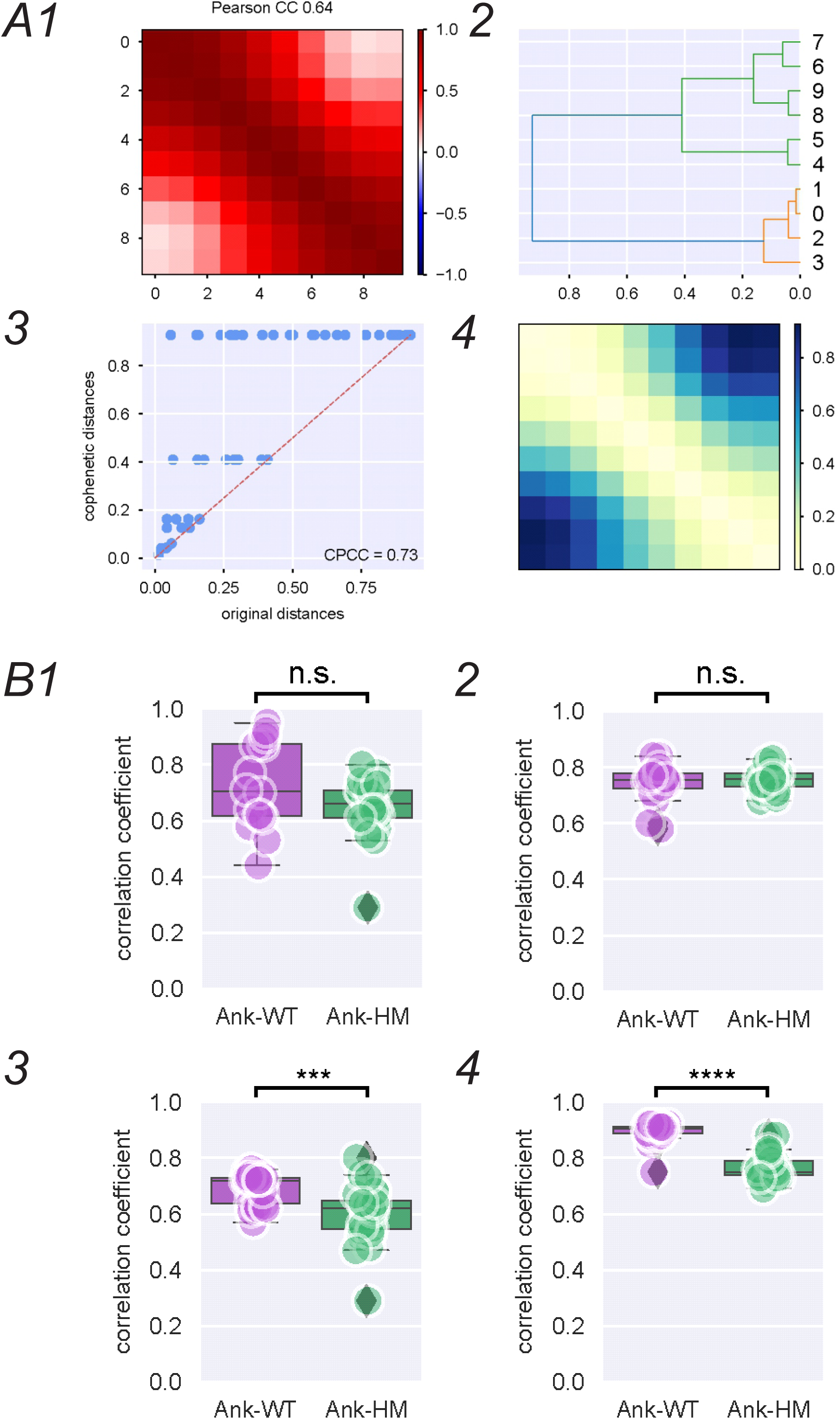
Impaired intra-columnar network synchronization in awake Ank3-1b^⁻/⁻^ mice. (A) Workflow for quantifying functional connectivity from laminar LFPs (100 ms post-stimulus). (A1) Representative cross-correlation matrix from an H.M. mouse. (A2) Hierarchical clustering (complete linkage) dendrogram of the correlation matrix from A1. (A3) Shepard plot for calculating the cophenetic correlation coefficient. (A4) Dissimilarity matrix (1 - Pearson correlation). (B) Group data for synchronization metrics. (B1, B2) Average Pearson correlation coefficient (B1) and cophenetic correlation coefficient (B2) under anesthesia. (B3, B4) Average Pearson correlation coefficient (B3, p < 0.001) and cophenetic correlation coefficient (B4, p < 0.001) in the awake state. Data are presented as box plots; ***p ≤ 0.001 (Mann-Whitney U-test).

Spike-field coherence (SFC) analysis compared cortical synchrony in response to air-puff stimuli between Ank3-1b^-/-^ and WT mice under anesthetized and awake conditions, focusing on activity within 500 ms from stimulus onset. Coherence was evaluated in two frequency bands: 3–30 Hz (low/mid frequencies) and 30–80 Hz (gamma range), based on power-spectral density (Fig. 6A). Ank3-1b^-/-^ mice exhibited stronger coherence in the low-frequency range (3–30 Hz) under both anesthetized (p < 0.05, Fig. 6B1) and awake conditions (p < 0.001, Fig. 6C1), but not in the gamma range (30–80 Hz; Fig. 6B2, C2). These findings suggest genotype- and state-dependent modulation of cortical oscillatory coupling in the low-frequency range, highlighting Ank3’s role in maintaining network synchrony during sensory processing. Surprisingly, deficits in parvalbumin interneuron excitability in Ank3-1b^-/-^ mice, known to critically contribute to gamma-band activity (Hadler, Tzilivaki et al. 2024), were not associated with differences in spike-field coherence (SFC) in the gamma band.

**Figure 6.**
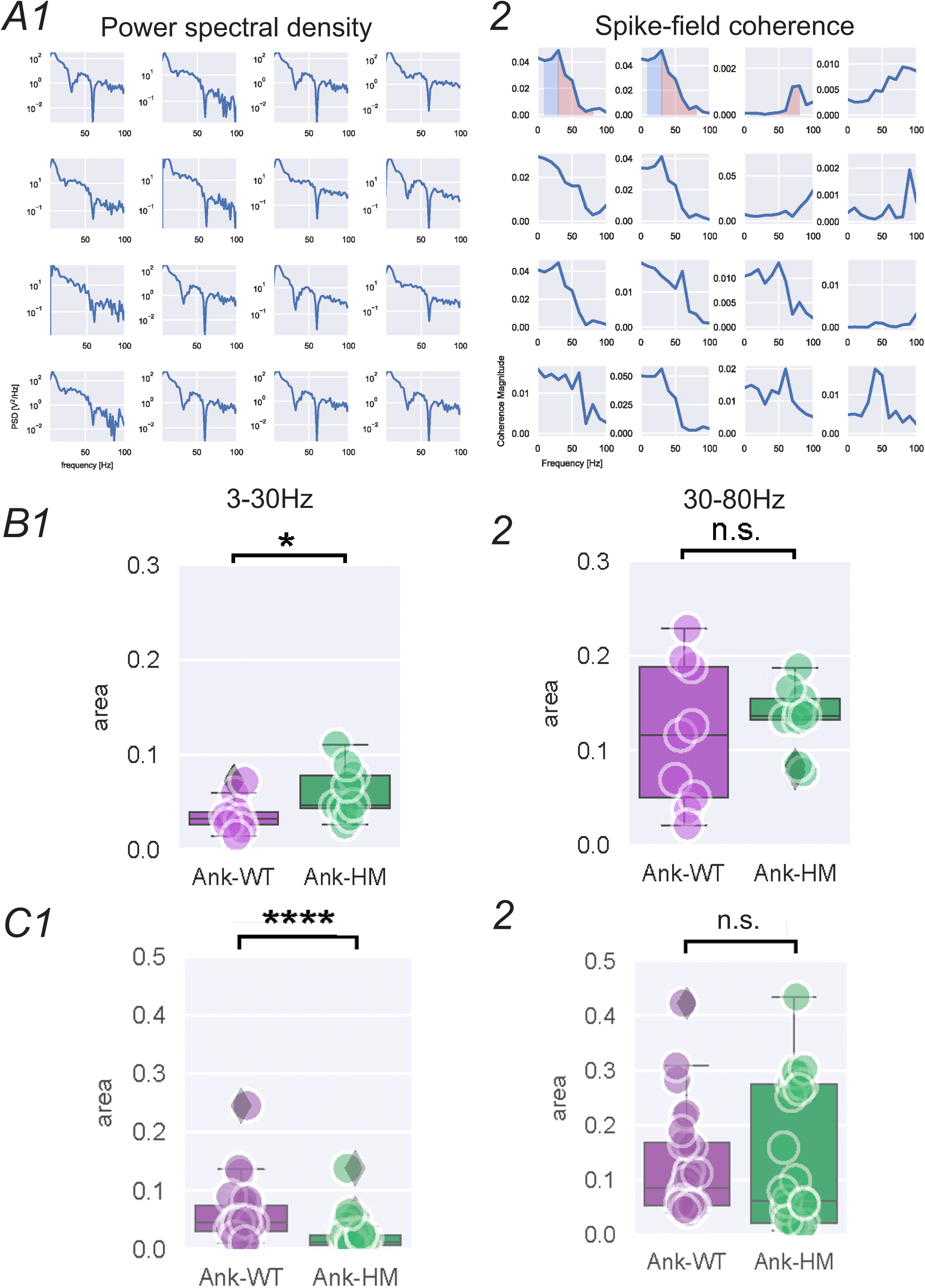
Altered low-frequency spike-field coherence (SFC) in Ank3-1b^⁻/⁻^ mice. (A1) An example of Power spectral density (PSD) of the LFPs from 16 channels under anesthetized conditions. (A2) Representative spike-field coherence plot from 16 channels, highlighting the low-frequency (3-30 Hz, blue) and gamma (30-80 Hz, red) bands. (B) Integral of spike-field coherence under isoflurane anesthesia for the 3-30 Hz band (B1, p < 0.05) and the 30-80 Hz band (B2, p = n.s.). Mann-Whitney U-test. (C) Spike-field coherence integrals across cortical channels in the awake state for the 3-30 Hz band C1, left) and the 30-80 Hz band (C2, right, 3-30 Hz: ****p ≤ 0.0001; 30-80 Hz: p = n.s., Mann-Whitney U-test).

### 3. Dynamics of Sensory-Evoked Excitatory and Inhibitory Single Units

In single-unit analyses comparing *Ank3-1b*^-/-^ and wild-type (WT) mice, putative excitatory (EXH) and inhibitory (INH) units were classified based on spike waveform features---trough-to-peak latency (time between the initial negative trough and subsequent positive peak) and asymmetry index (peak amplitude asymmetry)---which distinguish fast-spiking INH neurons (shorter trough-to-peak latency, symmetric waveforms) from regular-spiking EXH neurons (longer latency, asymmetric waveforms)(Reyes-Puerta, Yang et al. 2016). Under anesthetized (1.75% isoflurane) conditions, significant differences in spike latency (time to first action potential post-stimulus) were observed between genotypes, as well as INH (Fig 7C, p<0.01) but not EXH (Fig 7C). In infragranular layers (V/VI), *Ank3-1b*^-/-^ INH units, but not EXH units, exhibited prolonged latencies compared to WT (p < 0.001, rank sum test), with median latencies increasing by ∼25% (*Ank3-1b*^-/-^: 52 ± 6 ms vs. WT: 42 ± 6 ms, Fig 7E, p<0.05). This delay was not significant in supragranular layers II/III), where *Ank3-1b*^-/-^ INH latencies were similar to WT (Fig 7D). In awake conditions, we did not separate EXH vs. INH units. The putative latency in *Ank3-1b*^-/-^ mice were significantly prolonged in supragranular layers (p=0.005), however, it shows the opposite pattern in the infragranular layer (P=0.033, Fig 7G). Thus, the Ank3-1b^-/-^ mutation disrupts sensory-evoked inhibitory neuron dynamics, causing prolonged spike latencies in infragranular layers under anesthetized conditions and layer-specific latency alterations in awake states, suggesting impaired cortical circuit timing in epileptic encephalopathy. These effects, primarily on inhibitory neurons, may contribute to sensory and cognitive deficits by disrupting excitatory-inhibitory balance. These layer-specific deficits align with disrupted thalamocortical integration in Infragranular and supragranular zones, consistent with clinical EE studies reporting impaired sensory processing in disorders like Lennox-Gastaut syndrome(Arzimanoglou, French et al. 2009).

**Figure 7.**
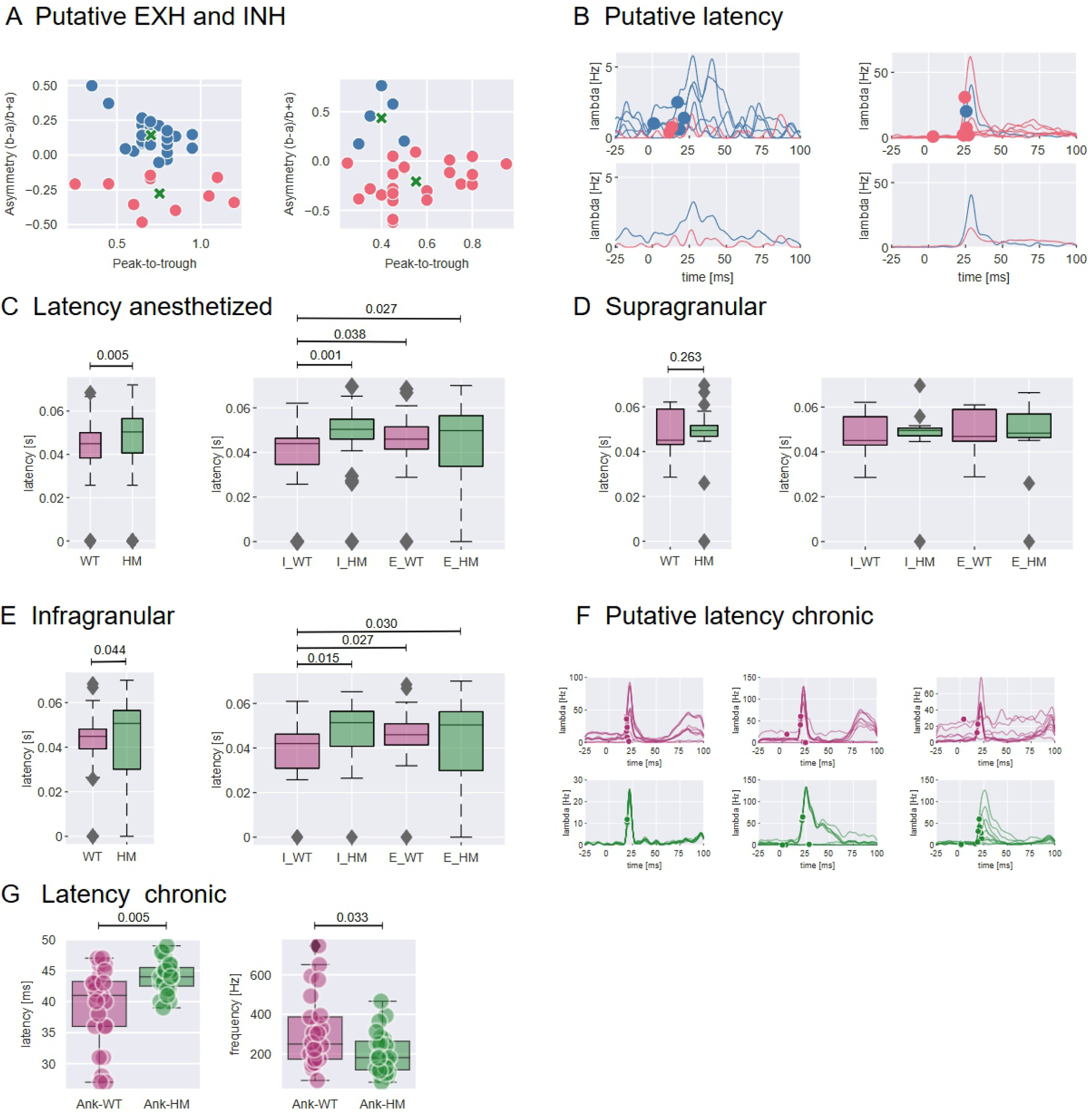
State- and layer-dependent disruption of sensory-evoked single-unit latencies. (A) Classification of putative excitatory (EXC, red) and inhibitory (INH, blue) neurons based on spike waveform features. (B) Representative spike density functions (SDFs) from a WT mouse, showing onset latencies (dots). (C) Population data of first-spike latencies under anesthesia (Mann-Whitney U-test). (D, E) Layer-specific latencies under anesthesia for supragranular (D) and infragranular (E) layers (rank-sum test). (F) Representative SDFs and latencies from awake recordings from WT (purple) versus mutant (green). (G) Latencies (p = 0.005) and frequencies (p = 0.033) in the awake state for all recordings pooled together (Mann-Whitney U-test). All data are presented as box plots.

### 4. Spike Adaptation Dynamics in Response to Repeated Sensory Inputs

To observe the cortical responses to repeated stimulations, we first performed spike density function (SDFs) estimation within each interval of different stimulations (1, 2 5 and 8 Hz air-puff). We observed significant adaptations (i.e. reduction SDF spikes under area in second-4^th^ air puff across four frequencies in both *Ank3-1b*^-/-^ and WT mice (Fig 8B), suggesting that spike-adaptation is preserved across genotypes. Mowery adaptation analysis(Mowery, Harrold and Alloway 2011) of cortical SDFs revealed that spike counts are negatively correlated with the stimulation frequencies in both genotypes (Fig 8B), and that there are no significant differences in the slope of this negative correlations (Fig 8C), suggesting preserved gross network output to repetitive whisker stimuli. Both Ank3-1b^-/-^ and WT mice exhibit significant spike adaptation to repeated whisker stimuli across all tested frequencies, with spike counts negatively correlated with stimulation frequency. The lack of significant differences in adaptation slopes between genotypes suggests that gross cortical network output in response to repetitive sensory stimuli remains preserved in Ank3-1b^-/-^ mice. These findings suggest that *Ank3-1b*^-/-^ mice cortices retain the capacity to integrate sensory inputs across diverse stimulation regimes, despite altered single-neuron excitability and disrupted temporal precision-likely reflecting deficient PV interneuron-mediated inhibition during early response phases.

**Figure 8.**
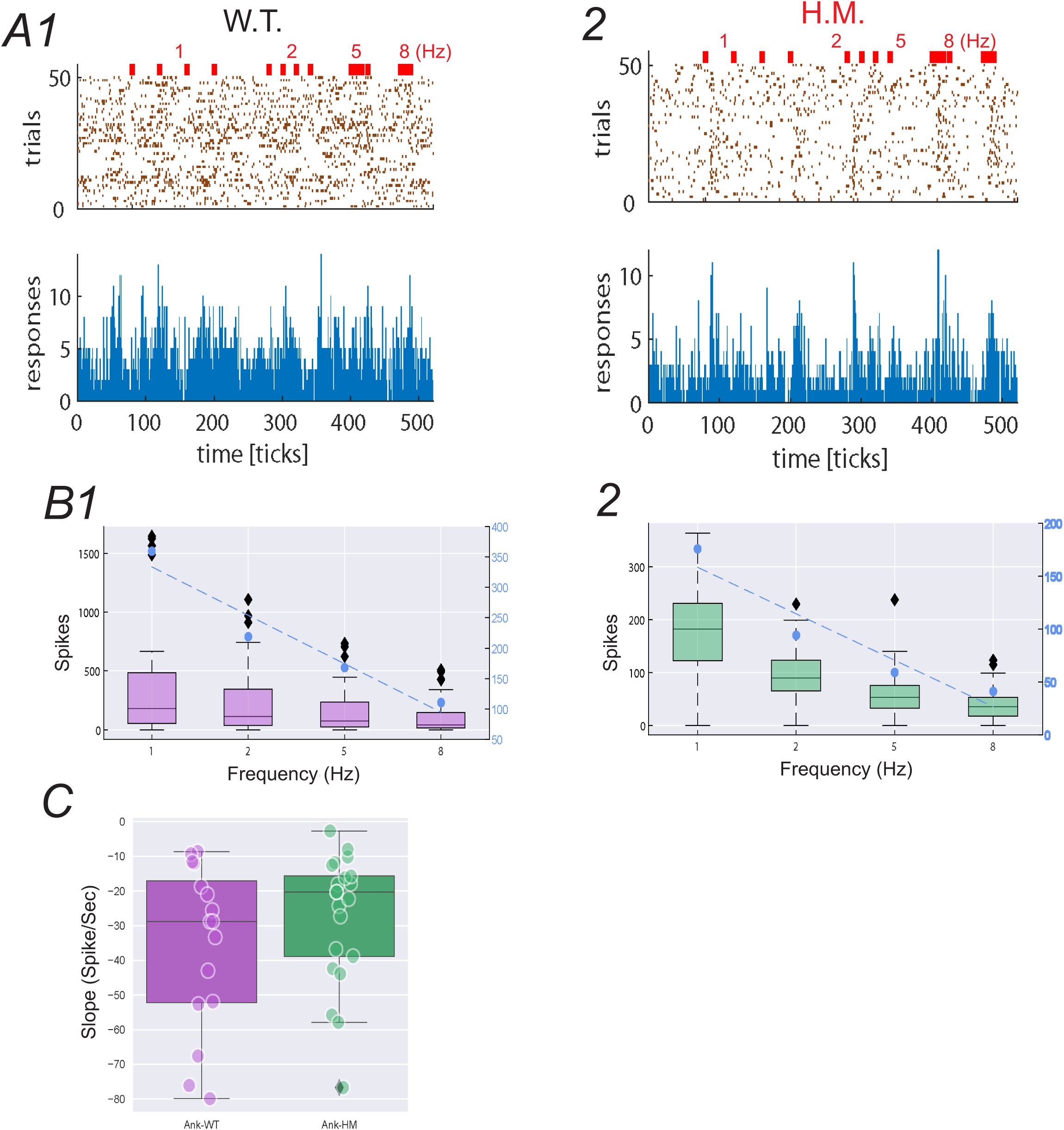
Preserved frequency-dependent adaptation of cortical sensory responses in Ank3-1b^⁻/⁻^ mice. (A) Representative raster plots and peri-stimulus time histograms (PSTHs, bin width = 25 ms) from sequential 1, 2, 5, and 8 Hz air puff stimulation in a WT (A1) and an H.M. (A2) mouse. (B) Box plots of the Spike Density Function (SDF) integral for each stimulus frequency in WT (B1) and H.M. (B2) mice. Average SDF integrals are shown (blue markers). (C) Slope of the linear fit to the average SDF integrals for each animal (p = n.s., Mann-Whitney U-test). Data in (B, C) are presented as box plots.

### 5. Sensory-Dependent Exploratory Behavior, Object Recognition, and Circadian Rhythm

In the open field test, Ank3-1b^-/-^ mice, but not WT, exhibited heightened anxiety-like behavior, spending 2.5 times (light on) and 3 times (light off) longer in the peripheral zone of the arena compared to the time spent in the center (Fig. 9B). Additionally, the absolute time spent in the peripheral zone was significantly longer in Ank3-1b^-/-^ mice compared to WT (Fig. 9B). Locomotor activity, measured as total distance traveled, did not differ between genotypes (p = 0.12), indicating that motor deficits in Ank3-1b^-/-^ mice did not confound the results. Thus, Ank3-1b^-/-^mice exhibit heightened anxiety-like behavior compared to WT mice, as evidenced by significantly increased time spent in the peripheral zone of the open field test under both light-on and light-off conditions. The absence of differences in locomotor activity between genotypes indicates that these behavioral differences are not confounded by motor deficits, suggesting that the Ank3-1b mutation specifically impairs anxiety-related sensory or emotional processing in this model of epileptic encephalopathy.

**Figure 9.**
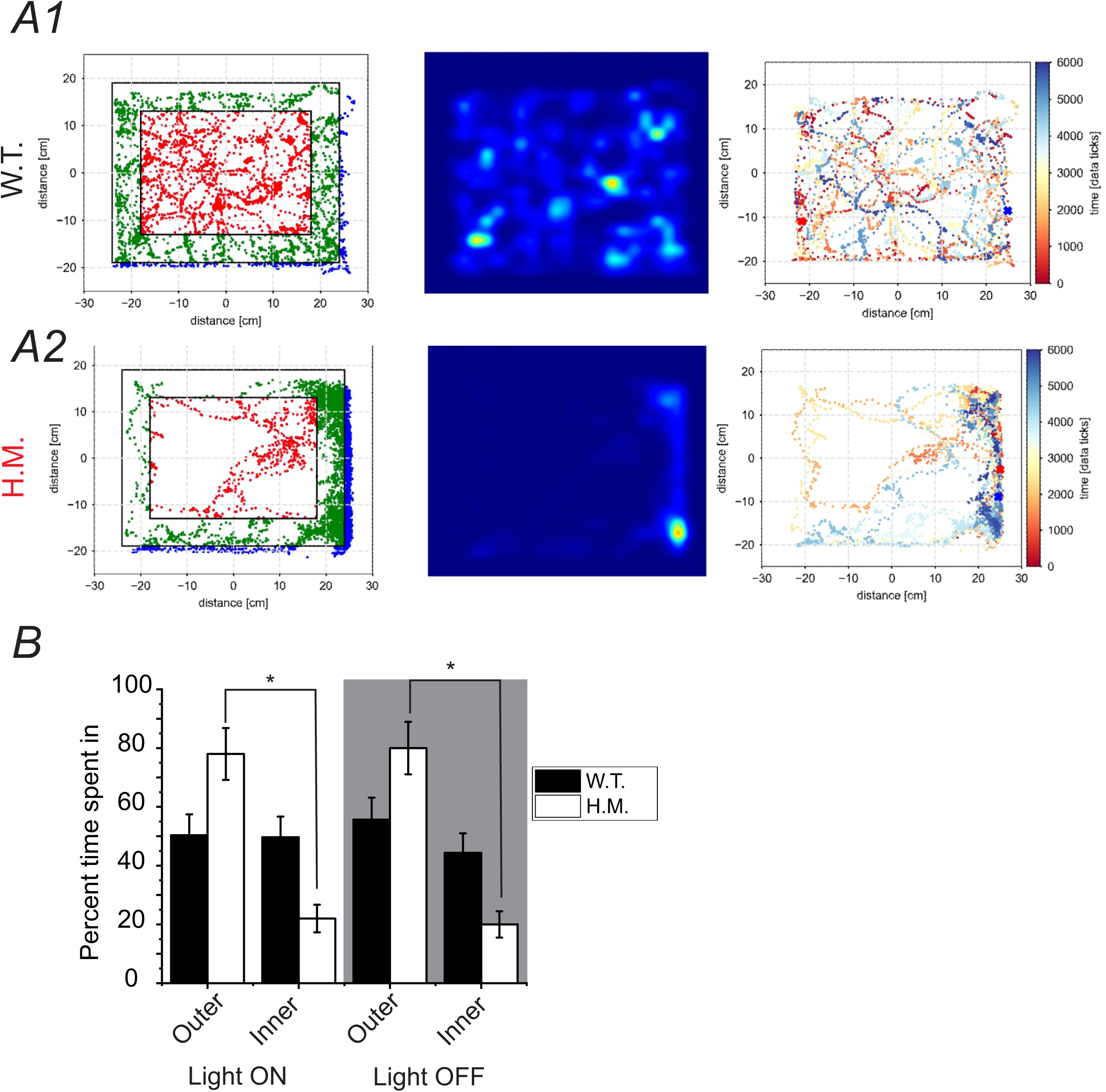
Ank3-1b^⁻/⁻^ mice exhibit anxiety-like behavior but intact recognition memory. (A) Representative locomotion tracks in the open field for a WT (A1) and an H.M. (A2) mouse. (B) Quantification of open field and novel object recognition tests. (B) Percent of time spent in the center versus the outer area of the open field under light and dark conditions.

Ank3-1b^-/-^ mice exhibited increased variance in the period length of daily locomotor activity (LMA, F_(6,6)_=276.3, p<0.0001) and core body temperature (Tb, F_(6,6)_=45.85, p=0.0002) rhythms compared to wild-type (WT) controls (Fig. 10A). While some Ank3-1b^-/-^ mice displayed period lengths outside the circadian range (22–26 h) for both LMA and Tb, others maintained periods within this range. The amplitude of LMA rhythms was reduced in mutants (t_(12)_=2.939, p=0.0124), and Tb rhythms showed greater amplitude variance (F_(6,6)_=15.86, p=0.0038) relative to WT mice (Fig. 10B). Additionally, Ank3-1b^-/-^ mice demonstrated altered phase angles of entrainment (defined as hours from lights-on to rhythm acrophase) for LMA rhythms, with increased variability in the timing of peak rhythmicity (F_(6,6)_=7.726, p<0.0252). Collectively, these observations suggest potential disruptions in circadian regulation in a subset of Ank3-1b^-/-^ mice, highly similar to diverse chronotype and its variability in epilepsy populations(Najar, Santos et al. 2024).

**Figure 10.**
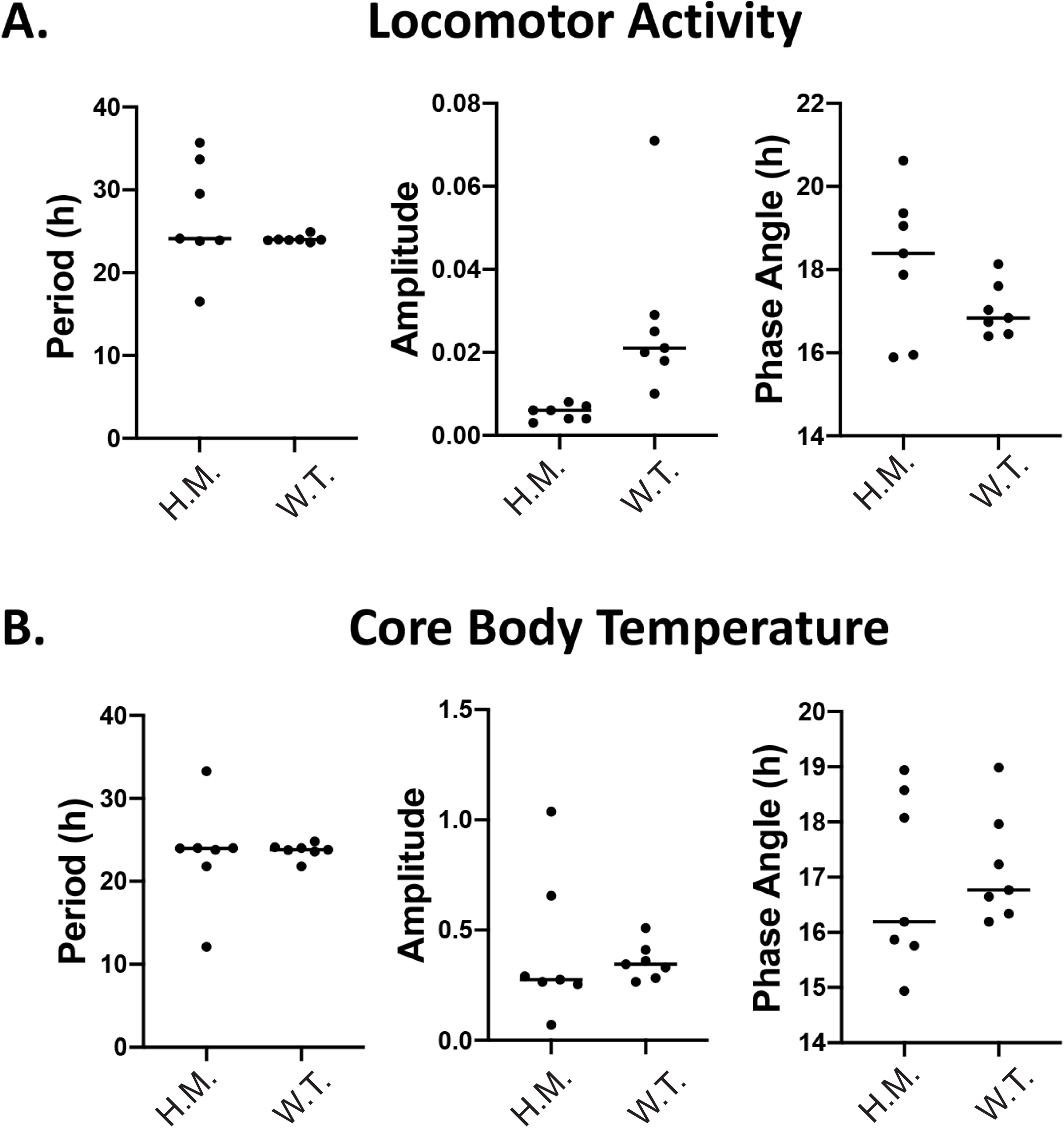
Circadian rhythm profiles of locomotor activity and core body temperature. Circadian group data from 7-day recordings of locomotor activity (A) and core body temperature (B) for H.M. and W.T. mice under a 12-hour light/dark cycle. Period (F_(6,6)_=276.3, p<0.0001), amplitude (t_(12)_=2.939, p=0.0124), and phase angle of entrainment for locomotor activity (F_(6,6)_=7.726, p<0.0252). Period (F_(6,6)_=45.85, p=0.0002), amplitude (F_(6,6)_=15.86, p=0.0038), and phase angle of entrainment for core body temperature. Data are presented as individual points with group median.

## Discussion

This study reveals that Ank3-1b^-/-^ mice, a genetic model of epileptic encephalopathy (EE) with parvalbumin (PV) interneuron dysfunction, exhibit profound cortical circuit abnormalities: disrupted excitatory-inhibitory (E-I) balance, laminar desynchronization, and delayed inhibitory signaling, while paradoxically preserving sensory adaptation, recognition memory, and select behavioral functions. These findings elucidate how ANK3 mutations disrupt cortical network dynamics and highlight compensatory mechanisms that mitigate behavioral deficits despite widespread circuit pathology. Below, we discuss the implications of these results across cortical dynamics, columnar sensory processing, behavioral outcomes, and circadian regulation, integrating cellular, network, and behavioral perspectives.

### Cortical Hyperexcitability and Laminar Desynchronization

Using laminar recordings and CSD analysis in rat barrel cortex, Di et al., found that vibrissa-evoked potentials consist of sequential fast and slow components(Di, Baumgartner and Barth 1990). The fast responses reflect sequential depolarization from supragranular to infragranular layers, while the slow waves represent subsequent repolarization processes. These findings indicate a basic cortical circuit of sequential pyramidal cell activation that generates evoked field potentials(Di, Baumgartner and Barth 1990). The reduced sink/source ratio (p = 0.0279) and amplified average rectified current (AVREC; p < 0.01) in Ank3-1b^-/-^ mice indicate cortical hyperexcitability, mirroring EEG hallmarks of EEs such as epileptiform discharges (McTague, Howell et al. 2016). This phenotype, driven by increased source magnitudes (454.8 ± 69.8 vs. 234.3 ± 60.6 µm²·s; p = 0.026) and altered sink areas (167.0 ± 29.8 vs. 289.5 ± 51.3 µm²·s; p = 0.034), suggests impaired coordination of excitatory and inhibitory currents, likely due to PV interneuron hypoexcitability (Lopez, Wang et al. 2016). Notably, AVREC amplification was state-dependent, peaking in awake conditions (0.0023 ± 0.0025 vs. 0.0008 ± 0.0015 µV/mm²; p < 0.0001) but not under anesthesia (p = 0.327), highlighting the role of brain state in modulating circuit dysfunction.

Cross-correlation analysis revealed reduced functional connectivity in awake Ank3-1b^-/-^ mice, with lower Pearson correlation coefficients (0.65 ± 0.21 vs. 0.76 ± 0.15; p < 0.001) and cophenetic clustering scores (0.78 ± 0.05 vs. 0.91 ± 0.09; p < 0.001) indicating desynchronization, particularly in supragranular layers (II/III) and layer IV. These findings align with impaired thalamocortical integration in EE syndromes like Lennox-Gastaut syndrome (Arzimanoglou, French et al. 2009) and suggest that Ank3-1b loss disrupts the temporal fidelity of cortical microcircuits during sensory processing. Surprisingly, spike-field coherence (SFC) showed enhanced low-frequency (3–30 Hz) coherence in Ank3-1b^-/-^ mice under both anesthetized (p < 0.05) and awake conditions (p < 0.001), but no differences in gamma-band (30–80 Hz) coherence, despite PV interneuron deficits known to drive gamma oscillations (Cardin 2018, Hadler, Tzilivaki et al. 2024). This dissociation suggests that low-frequency oscillations may dominate pathological network states in EEs, potentially disrupting higher-frequency synchrony(Lenck-Santini 2017).

### Columnar Processing of Sensory Inputs

The layer-specific deficits in Ank3-1b^-/-^ mice provide insights into columnar sensory processing. Prolonged inhibitory unit latencies in infragranular layers (V/VI; 52 ± 6 ms vs. 42 ± 6 ms; p < 0.001) but not supragranular layers (II/III) under anesthesia indicate selective disruption of deep-layer inhibitory circuits, which are critical for feedback modulation of sensory inputs (Douglas and Martin 2004). In awake conditions, the reversal of latency patterns—prolonged in supragranular layers (p = 0.005) but reduced in infragranular layers (p = 0.033)—suggests state-dependent compensatory mechanisms or altered thalamocortical drive. These findings extend prior work on Ank3-1b^-/-^ mice (Lopez, Wang et al. 2016) by demonstrating that AnkG loss at PV interneuron axonal initial segments (AIS) impairs the temporal precision of columnar processing, potentially contributing to sensory hypersensitivity observed in EE patients (Zuberi, Wirrell et al. 2024).

Despite these deficits, Ank3-1b^-/-^ mice exhibited preserved sensory adaptation to repeated whisker stimuli (1–8 Hz), with comparable spike density function (SDF) integrals and Mowery adaptation slopes across genotypes (Fig. 8B–D). This resilience suggests that cortical networks prioritize maintaining gross sensory output over fine temporal coding, possibly through redundant excitatory pathways or compensatory interneuron types (e.g., somatostatin interneurons; (Nolan, Sohal and Rosi 2022). The intact adaptation contrasts with Scn1a haploinsufficient mice, where sensory deficits are pronounced(Favero, Sotuyo et al. 2018), highlighting distinct roles of Ank3 and Scn1a in sensory circuit organization.

### Behavioral Dissociations and Compensatory Mechanisms

The behavioral profile of Ank3-1b-/- mice reveals a striking dissociation between circuit pathology and function(Berg, Berkovic et al. 2010). Heightened anxiety-like behavior in the open field test (2.5–3 times longer in peripheral zones; p < 0.05) aligns with EE comorbidities and may reflect amygdala hyperactivity secondary to cortical desynchronization (Shao and Stafstrom 2016) (Pellock and Brittain 2016). In contrast, the preserved sensory functions despite E-I imbalance and desynchronization suggest compensatory mechanisms, such as redundant cortical wiring or plasticity in non-PV interneuron populations. The intact sensory adaptation further indicates that Ank3-1b-/- circuits prioritize functional output over precise timing, unlike other EE models (e.g., Kcnq2; (Bender, Natola et al. 2013)).

### Entrained Circadian Rhythm Variability

Ank3-1b^-/-^ mice exhibited increased variability in locomotor activity (LMA) and core body temperature (Tb) rhythms under entrained conditions (12h/12h light-dark), with some showing period lengths outside the circadian range (22–26 h), reduced LMA amplitude, and altered phase angle of entrainment (Fig. 10A–B). These differences suggest potential circadian dysregulation in a subset of mutants, consistent with disrupted sleep-wake cycles in some EE patients (Sanchez Fernandez, Gainza-Lein et al. 2019). These results align with ANK3’s role in regulating neuronal excitability.

### Mechanistic and Therapeutic Implications

Our findings establish Ank3-1b^-/-^ mice as a robust model for studying EE circuit mechanisms, linking AnkG loss to laminar-specific desynchronization, delayed inhibition, and hyperexcitability. The selective impact on low-frequency coherence and preserved gamma-band SFC despite PV deficits suggest that Ank3 modulates oscillatory coupling indirectly, possibly via altered NaV channel clustering or AIS integrity (Rasband 2010). These insights have therapeutic implications: targeting low-frequency oscillations or enhancing PV interneuron function (e.g., via optogenetic stimulation or pharmacological modulators) could restore temporal precision and mitigate sensory and anxiety phenotypes.

### Limitations and Future Directions

Several limitations warrant consideration. First, the NOR task’s reliance on exploratory drive may have obscured subtle cognitive deficits, particularly given the heightened anxiety in Ank3-1b^-/-^ mice. Future studies should employ tactile discrimination or working memory tasks to probe cognition more robustly. Second, the lack of excitatory vs. inhibitory unit classification in awake conditions limits mechanistic specificity; optogenetic tagging could address this. Third, the non-significant circadian findings suggest a need for larger sample sizes or longitudinal recordings to clarify Ank3’s role in circadian regulation. Finally, exploring compensatory roles of non-PV interneurons (e.g., somatostatin or VIP) or glial cells could elucidate resilience mechanisms.

## Conclusion

Ank3-1b^-/-^ mice exhibit profound cortical circuit dysfunction-hyperexcitability, desynchronization, and delayed inhibitory signaling-yet maintain sensory adaptation and recognition memory, likely due to compensatory cortical wiring. These findings highlight the complex interplay between circuit pathology and behavior in EEs, underscoring the need for therapies targeting temporal precision and low-frequency oscillations. By linking ANK3 mutations to disrupted columnar processing and network synchrony, this study provides a framework for developing circuit-based interventions to address EE and its comorbidities.

## Materials and Methods

## 1. Animals

Male and female Ank3-1b^-/-^ mice (Lopez, Wang et al. 2016) and wild-type (WT) littermates (postnatal days P30–P90) were bred from heterozygous Ank3-1b+/- parents on a C57BL/6J background. Mice were group-housed (4–5 per cage) under a 12-hour light/dark cycle (lights on at 07:00) with ad libitum access to food and water. Genotyping was performed via PCR on tail biopsies using primers specific for Ank3 exon 1b (forward: 5′-CAGGTGTTGGGTCAGTTTCC-3′; reverse: 5′-GCCACAGTCATAGCAGTGGT-3′) as described(Lopez, Wang et al. 2016). All procedures were approved by the University of Wyoming Institutional Animal Care and Use Committee (IACUC) and adhered to NIH guidelines.

## 2. Surgical Procedures

### 2.1. Head-Plate Implantation and Chronic Probe Insertion

Mice were anesthetized with isoflurane (5% induction, 1.5–2% maintenance in 100% O₂) and secured in a stereotaxic frame (Kopf Instruments). Body temperature was maintained at 37°C using a heating pad. A custom titanium head-plate (5 × 5 mm) was affixed to the skull with dental cement (C&B-Metabond) over a 2 × 2 mm craniotomy centered on the right primary somatosensory barrel cortex (S1BF; AP: −1.82 mm, ML: +3.25 mm from bregma). A 16-channel linear silicon probe (A1×16-3mm-100-177, NeuroNexus; 100 µm inter-electrode spacing) was implanted at a 30° angle to a depth of 3.3 mm using a hydraulic micromanipulator (Narishige). The craniotomy was sealed with Kwik-Cast (World Precision Instruments), and the probe was secured with dental cement. A silver wire reference electrode was placed in the olfactory bulb (AP: +4.0 mm, ML: 0 mm). Postoperative care included buprenorphine (0.1 mg/kg, subcutaneous) and ibuprofen (50 mg/kg, intraperitoneal) for 72 hours, with daily monitoring for weight and behavior.

### 2.2. Neurotar Mobile HomeCage Training

To prepare for awake recordings, mice underwent habituation to the Mobile HomeCage® (MHC; Neurotar) as follows: Pre-surgical: Mice were acclimated to handling via tunnel transfer (5 cm diameter PVC tube) for 7 days (10 min/day). Post-surgical: Mice were gradually exposed to the MHC air table (30–60 min/day) over 4 days. Head-fixation training progressed from 2.5 to 30 min/day, with mice navigating a carbon-fiber tray via airflow propulsion to ensure comfort during recordings.

### 2.3. Biotelemetry assessment of circadian function

For biotelemetry transmitter implantations, mice were anesthetized with a ketamine (100 mg/kg) and xylazine (10 mg/kg) mixture diluted 1:10 in saline, via intraperitoneal injection. The belly was shaved and sterilized with alcohol and iodine prep pads and a midline incision was made (1 cm). A biotelemetry transmitter (TAF10, Data Sciences International) was inserted into the intraperitoneal cavity, and the internal membrane and external skin were closed with sutures. Mice were given buprenorphine (0.1 mg/kg) in saline subcutaneously for postoperative analgesia followed by 1 mL of saline to account for fluid loss.

## 3. Electrophysiological Recordings

Recordings targeted the S1 barrel cortex to assess sensory-evoked responses to whisker stimulation under anesthetized and awake conditions. Whisker stimuli were delivered via air-puffs (20 psi, 50 ms duration, 0.1–8 Hz, PulsePal) using a home-made solenoid valve connected to pressured airs.

### 3.1. Anesthetized Recordings Mice were anesthetized with isoflurane (1–1.75% in 100% O₂) and placed in a stereotaxic frame

A 16-channel linear probe (A1×16-3mm-100-177; impedance: 1–1.5 MΩ) or a 32-channel linear Edge probe (A1×32-Edge-5mm-100-177; NeuroNexus) was inserted into S1BF. Local field potentials (LFPs) and multi-unit activity were amplified (1000×), bandpass-filtered (0.1–7500 Hz), digitized at 20 kHz (SmartBox 2.0, NeuroNexus), and notch-filtered (60 Hz) to remove line noise.

### 3.2. Awake Recordings. Head-fixed mice in the MHC underwent extracellular recordings during whisker stimulation

Signals were acquired at 30 kHz using an RHD2000 system (Intan Technologies) with identical filtering. Recording protocols included: Spontaneous Activity: 10 min baseline to assess intrinsic network activity. Sensory-Evoked Responses: 200 trials of 0.5 Hz air-puff stimulation for current source density (CSD) and single-unit analysis. Frequency Adaptation: 50 trials each at 1, 2, 5, and 8 Hz to assess Mowery adaptation.

## 4. Behavioral Testing

Behavioral assays were conducted in a sound-attenuated room under dim red light (10 lux) unless specified. Mice were habituated to the testing room (30 min/day) for 3 days prior to experiments. All sessions were recorded and analyzed using Ethovision XT® v15 (Noldus Information Technology).

### 4.1. Open Field (OF) Test

Mice freely explored a 40 × 40 × 40 cm acrylic arena for 10 min under light-on (100 lux) and light-off (0 lux) conditions, tested on separate days. A 25 × 25 cm central zone was defined as the “center.” Total distance traveled, velocity, and time spent in the center vs. periphery were quantified to assess locomotor activity and anxiety-like behavior (Kalueff, Wheaton and Murphy 2007).

### 4.2. Novel Object Recognition (NOR) Test. Mice explored two identical objects (plastic cylinders, 5 cm height) in a 40 × 40 cm arena for 10 min (Phase 1)

After a 24-hour delay, one object was replaced with a novel object (wooden cube, 5 cm) for 5 min (Phase 2). Exploration was defined as snout within 2 cm of an object, excluding climbing or rearing. Discrimination index (DI) and recognition index (RI) were calculated as:

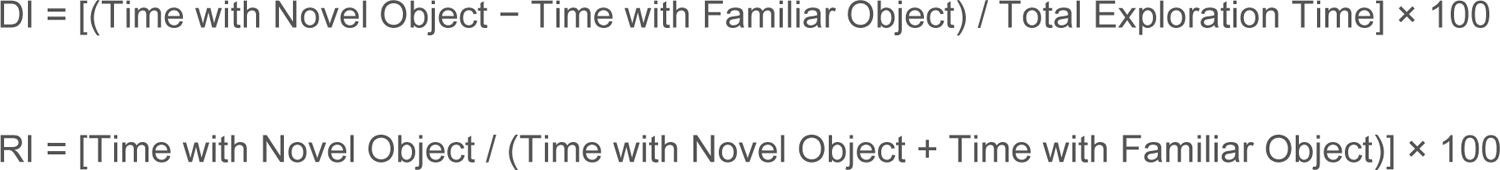

### 4.3. Circadian Rhythm Monitoring

After one week recovery from telemetry transmitter implantation surgery, mice were moved in their home cages into light-tight and noise-attenuated circadian chambers (Phenome) and given at least five days to acclimate before recordings began. Home cages sat atop telemetry receivers connected to a computer and data acquisition system (Data Sciences International, Ponemah software). Clocklab Chamber Control software (Actimetrics) was used to program the light-dark cycle (12h-ON, 12h-OFF) and to measure light levels (300lux during the light period) as well as ambient temperature (23.5±1°C) and humidity levels (25-32%). Locomotor activity (LMA) and core body temperature (Tb) were monitored using these implanted transmitters under a 12-hour light/dark cycle, and data were collected continuously for 7 consecutive days. Period length, amplitude, and phase angles of entrainment (hours from lights-on to rhythm acrophase) were analyzed using ClockLab (Actimetrics) to assess circadian rhythmicity (Jud, Schmutz et al. 2005).

## 5. Histology

Mice were anesthetized with ketamine/xylazine (100/10 mg/kg, intraperitoneal) and transcardially perfused with 4% paraformaldehyde in 0.1 M phosphate buffer (pH 7.4). Brains were cryoprotected in 30% sucrose, sectioned coronally at 40 µm (Leica CM3050 S cryostat), and stained with cresyl violet (Nissl) or DAPI (Vectashield). Cytochrome oxidase (CO) staining was performed to identify S1BF barrels: Free-floating sections were rinsed in 0.1 M phosphate buffer (pH 7.4). Sections were incubated in a reaction solution (0.05% cytochrome C, 0.02% catalase, 0.05% diaminobenzidine, 10% sucrose in 0.1 M phosphate buffer) at 37°C for 2–4 hours. Reactions were stopped by rinsing in buffer, followed by dehydration in graded ethanol, clearing in xylene, and coverslipping with DPX. Barrel fields were visualized under brightfield microscopy (Zeiss Axio Imager 2) based on CO-dense regions. Electrode tracks were confirmed using DiI (2 mg/ml in ethanol) applied to probes before implantation, imaged with fluorescence microscopy (Zeiss), and aligned with CO-stained sections to verify S1BF targeting.

## 6. Data Analysis

### 6.1. Current Source Density (CSD)

LFPs were re-referenced to a common average, down-sampled to 1 kHz, and analyzed using the standard CSD method(Nicholson and Freeman 1975): 1) 5-point approximation of the second spatial derivative:

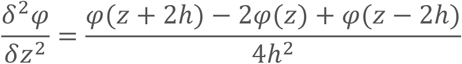

Then 1-dimensional CSD was computed in the following manner:

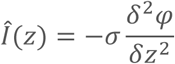

where Î(*Z*) is the current source density, φ is the LFP, *Z* is the depth, *h* is the inter-contact spacing, and σ is the tissue conductivity. Sink/source maps and average rectified current (AVREC) were generated using custom Python scripts. Sink/source areas and ratios were quantified over 500 ms post-stimulus.

### 6.2. Single-Unit Analysis

Spikes were detected using a threshold-based algorithm (4× RMS noise) and sorted with Kilosort4 (Pachitariu, Sridhar et al. 2024). Putative excitatory and inhibitory units were classified based on spike waveform features: Trough-to-peak latency (<0.5 ms: inhibitory; >0.5 ms: excitatory)(Reyes-Puerta, Yang et al. 2016). Asymmetry index (peak amplitude ratio). Peri-stimulus time histograms (PSTHs; 5 ms bins) and spike latencies (time to first action potential post-stimulus) were computed using NeuroExplorer (Nex Technologies).

### 6.3. Spike-Field Coherence (SFC)

Spike-Field Coherence (SFC) was computed to quantify the phase-locking between the timing of individual neuron action potentials (spikes) and the ongoing oscillations of the local field potential (LFP) within the cortical column. All analyses were performed on data from the 500 ms post-stimulus epoch to focus on stimulus-evoked neural synchronization. SFC was calculated using the built-in coherence functions in NeuroExplorer software (version 5, Nex Technologies).

#### Spectral Estimation and Coherence Calculation

For each single unit and the LFP recorded from the same microelectrode, the spike train was represented as a binary point process, and the LFP was treated as a continuous analog signal. The coherence, *C*_*s*f_(*f*), which measures the linear relationship between the two signals as a function of frequency *f*, was estimated using the Magnitude-Squared Coherence formula:

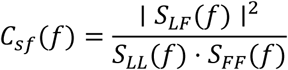

where:

- *S*_*LL*_(*f*) is the power spectral density of the LFP signal.
- *S*_*FF*_ (*f*) is the power spectral density of the spike train (estimated as the spectrum of a point process).
- *S*_*LF*_(*f*) is the cross-spectral density between the LFP and the spike train.

To ensure robust, low-variance spectral estimates, we employed a multi-taper method(Mitra and Pesaran 1999, Jarvis and Mitra 2001). Specifically, the Thomson multi-taper approach was used with **5 discrete prolate spheroidal (Slepian) tapers** and a **time-bandwidth product of 3**. This configuration provides an optimal balance between spectral resolution and estimation variance, effectively reducing noise in the coherence estimate while controlling for spectral leakage.

#### Frequency Band Analysis

The computed SFC was evaluated within two physiologically relevant frequency ranges:

1. **Low/Mid Frequency Band (3–30 Hz):** Encompassing theta (3-8 Hz), alpha (8-12 Hz), and beta (12-30 Hz) oscillations, often associated with long-range communication and top-down processing.
2. **Gamma Frequency Band (30–80 Hz):** Associated with local circuit processing, feature binding, and bottom-up sensory encoding.

For statistical comparisons, the mean coherence value within each of these defined bands was extracted for each unit-LFP pair.

### 6.4. Sensory Adaptation Analysis

The adaptation of neural responses to repetitive sensory stimuli was quantified using a protocol and analytical approach adapted from Mowery, Harrold et al.(Mowery, Harrold and Alloway 2011) (2011). Neuronal activity was recorded in response to a train of air-puff stimuli delivered at frequencies of 1, 2, 4, and 8 Hz, mimicking the detection of varying tactile flutter.

#### Spike Density Function (SDF) and Response Quantification

For each isolated single unit, peri-stimulus time histograms (PSTHs) were constructed for each stimulus frequency. To generate a continuous representation of instantaneous firing rate, PSTHs were converted into Spike Density Functions (SDFs) by convolving the spike train with a Gaussian kernel (σ = 200 ms). The Spike Density Function SDF(*t*) is generated by convolving a spike train, represented as a sum of Dirac delta functions δ(*t* − *t*_*i*_), with a Gaussian kernel *g*(*t*).

The formal equation is:

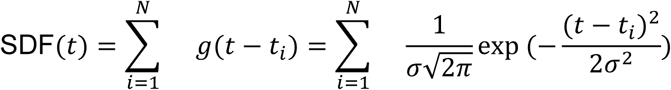

Where:

- SDF(*t*) is the estimated instantaneous firing rate at time *tt*.
- *N* is the total number of spikes in the spike train.
- *t*_*i*_ is the time of the *i*-th spike.
- *g*(*t*) is the Gaussian kernel function.
- σ is the standard deviation of the Gaussian kernel, which determines the temporal smoothing width (in your case, **σ = 200 ms**).
- 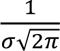 is the normalization constant for the Gaussian distribution.

The neural response to each stimulus in the train was quantified by calculating the integral of the SDF over a 500 ms post-stimulus window for each stimulus presentation. This integral is proportional to the total spike count evoked by the stimulus. For each frequency, the response magnitude was taken as the mean integral value across all stimuli within the train.

#### Adaptation Slope Calculation

To derive a single metric representing the rate of adaptation, a linear regression was performed on the mean response magnitudes across the four stimulus frequencies. The stimulus frequencies (1, 2, 4, 8 Hz) were log₂-transformed for the regression to linearize the exponential frequency scale, resulting in the predictor variable: Log₂(Frequency). The dependent variable was the corresponding SDF integral (spike count). The slope of the linear regression line, termed the **“Adaptation Slope,”** was used for subsequent statistical comparisons. A more negative slope indicates a steeper decline in response magnitude with increasing stimulus frequency, signifying stronger adaptation.

### 6.5. Functional Connectivity and Network Modularity Analysis

To assess functional connectivity and the hierarchical organization of cortical microcircuits, we calculated pairwise Pearson correlation coefficients followed by hierarchical clustering and cophenetic correlation analysis on local field potential (LFP) data.

#### 6.5.1. Cross-Correlation Matrix Construction

LFP traces from all recording channels, spanning a 500 ms post-stimulus window, were used without additional band-pass filtering. For each trial, pairwise functional connectivity was quantified by computing the Pearson correlation coefficient (PCC) between all possible pairs of recording channels *X* and *Y*. The PCC, a measure of linear covariation between two signals, was calculated as:

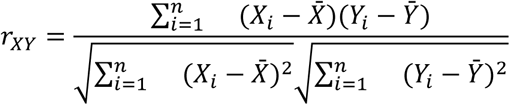

where *X*_*i*_ and *Y*_*i*_ are the voltage samples at time point *i* for the two channels, *X^-^* and *Y^-^* are the mean voltages of the respective signals over the 500 ms epoch, and *n* is the total number of samples in the epoch (Zar JH, 2010). The resulting *r*_*XY*_ values for all channel pairs were assembled into a single cross-correlation matrix per trial. Trial-averaged cross-correlation matrices were then computed for each animal and experimental condition.

#### 6.5.2. Hierarchical Clustering and Cophenetic Correlation

To evaluate the modular organization of the cortical network, we performed hierarchical clustering on the trial-averaged cross-correlation matrices using the unweighted average distance (UPGMA) linkage algorithm(Rohlf, Chang et al. 1990). The input to the clustering algorithm was a dissimilarity matrix, *D*_orig_, derived from the cross-correlation matrix, where the dissimilarity between channels *X* and *Y* was defined as *d*_orig_ = 1 − *r*_*X*Y_.

The output of hierarchical clustering is a dendrogram, a tree structure that illustrates the arrangement of the clusters. The faithfulness of this dendrogram in representing the original pairwise dissimilarities in *D*_orig_ was quantified using the cophenetic correlation coefficient (CCC) (Rohlf, Chang et al. 1990). For this, a cophenetic distance matrix, *D*_coph_, was constructed from the dendrogram, where the cophenetic distance between two channels is the height of the dendrogram node at which those two channels are first merged into a common cluster. The CCC was then calculated as the Pearson correlation coefficient between the elements of the original dissimilarity matrix *D*_orig_ and the cophenetic distance matrix *D*_cop*h*_:

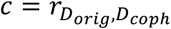

A CCC value close to +1 indicates that the dendrogram is a highly accurate representation of the original data structure, implying strong modularity, whereas lower values indicate a less faithful representation and disrupted network organization [3]. Statistical comparison of CCC values between groups was performed using the non-parametric Mann-Whitney U-test.

### 6.6. Circadian Analysis

LMA and Tb rhythms (7 consecutive days) were analyzed using ClockLab software. Period lengths were determined via chi-square periodogram, amplitudes (peak-to-mean) via cosinor analysis, and phase angles via cosine fitting (Jud et al., 2005). Mean differences were assessed using unpaired two-tailed t-tests and variance was assessed using F tests.

### 6.7. Statistics

Data are presented as mean ± SEM. Normality was tested using Shapiro-Wilk. Parametric tests (Student’s t-test, unpaired; mixed-effects ANOVA with Tukey’s post-hoc) were used for normally distributed data; non-parametric tests (Mann-Whitney U, Holm-Sidak post-hoc) were used otherwise. Analyses were performed in MATLAB (MathWorks), OriginPro (OriginLab), Prism 10 (GraphPad), or NeuroExplorer. Significance was set at p < 0.05.

## Acknowledgements

We thank C. Zhang for animal husbandry, animal genotyping, histology assistance and items purchasing. This work is supported by grants from National Institute of Mental Health (1R21MH131363), National Institute of Biomedical Imaging and Bioengineering (1R21EB032609), National Institute on Aging (5R21AG072803) and from National Institute of General Medical Sciences (2P20GM121310).

## Author contributions

Q.Q.S. designed the experiment and supervised the project, acquired the funding, and wrote the manuscript. D.S. co-designed the experiments, collected and analyzed data. T.C. performed statistical analysis. W.T performed circadian experiments, analyzed the data.

